# Identification and characterisation *of Mansonella perstans* in the Volta Region of Ghana

**DOI:** 10.1101/2023.11.15.567307

**Authors:** Millicent Opoku, Dziedzom K. de Souza

## Abstract

*Mansonella spp*. have been reported to have a wide global distribution. Despite the distribution and occurrence with other filarial parasites like *Wuchereria bancrofti, Onchocerca volvulus* and *Lao loa*, it is given little attention. Hence, there is no surveillance programme for assessing the distribution or mansonellosis, with mild to no symptoms experienced by infected people. However, addressing this infection is critical to the onchocerciasis control program as current rapid diagnostic tools targeting *O. volvulus* have the tendency to cross react with *Mansonella species*. In this study we identified and characterised *M. perstans* from five sites in the Volta region of Ghana and compared them to samples from other regions. Night blood smears and filter blood blots were obtained from individuals as part of a study on lymphatic filariasis. The giemsa stained smears were screened by microscopy for the presence of filarial parasites. Genomic DNA was extracted from blood blots from 39 individuals that were positive for *M. perstans* and Nested PCR targeting the internal spacer 1 (ITS-1) was conducted. Of these, 30 were sequenced and 24 sequences were kept for further analysis. Phylogenetic analysis of 194 nucleotide positions showed no differences in the samples collected. Samples from Ghana clustered with samples from other reference sequences from Africa and Brazil, possibly supporting the introduction of *M. perstans* from sub-Saharan Africa, during the trans-Atlantic slave trade. This study draws further attention to a neglected infection and presents the first characterisation of *M. perstans* in Ghana and calls for more population-based studies across different geographical zones to ascertain species variations and disease distribution.

## Introduction

Mansonellosis is known to occur in the tropics and subtropics with most of the cases in sub–Saharan Africa, some part Latin America, Brazil and the Caribbean. Overall, about 114 million people in Africa are estimated to be infected (1) and close to 600 million people are at risk (2). It is caused by the filarial nematodes, *Mansonella perstans, M. ozzardi* and *M. streptocerca*. Transmission is through the bite of infective midges (*Culicoides*) and *Simulium* blackflies (transmitting *M. ozzardi* in parts of Latin America) (3-6). Some mosquito species including Aedes and Anopheles have been implicated (4, 7, 8). *M. ozzardi* are the major species in Latin America and the Caribbean, while *M. perstans* and *M. streptocerca* are reported in Africa. *M. perstans* have also been reported along the coastal areas of equatorial Brazil and the Caribbean. Symptoms of mansonellosis range from fever, headache, pruritus, eosinophilia, to pain in bursae, joint synovia, in serous cavities and abdominal pain, proptosis, swelling of the eye and nodules in the conjunctiva (9-11). Diagnosis is by demonstration of microfilariae (which shows no periodicity) in peripheral blood (12). Drugs including diethylcarbamazine, ivermectin, mebendazole and doxycycline (which targets the bacteria endosymbiont Wolbachia) have been used to successfully clear the adult worms and reduce microfilariae burden in humans (13-16).

Although it occurs with other disease-causing filarial parasites such as lymphatic filariasis, loaisis and onchocerciasis, and has a wide distribution, symptoms of *Mansonella* infections are often mild and asymptomatic hence, often neglected (17). Hence, it receives little attention and only detected during structured surveillance for lymphatic filariasis and onchocerciasis (18). It is not considered to be of public health importance, as such no surveillance and management activities or funding are available to assess the burden and implement control activities (1, 3).

It is however, believed that the distribution of ivermectin through MDA campaigns for lymphatic filariasis (LF) and onchocerciasis may have impacted *Mansonella* infection prevalence (19). The evidence presented by some studies, indicating that 10% of all infections may lead to disease morbidity (17), calls for further studies in understanding the health impact of this widely distributed filarial parasite. Furthermore, the occurrence with other filarial parasites poses a great challenge to the development of new diagnostics. For instance, crossreactivity with serology-based rapid diagnostic test (RDT) for LF often leads to over estimations of LF prevalence, posing challenges to evaluating intervention efforts (13, 20-22).

Mansonellosis was first reported in the 1960s in Ghana. However, its pathogenicity is poorly understood. Few studies have tried to ascertain the epidemiology, with a paucity of information. Following on from 1960, a report on coinfection with Buruli Ulcer was reported in a community in the middle belt of Ghana (23). Subsequently, Debra and colleagues also reported an average of 32% infection in communities in the middle belt of Ghana (24). There hasn’t been any extensive work in any other parts of the country to examine the distribution, nor establish its pathogenicity. This study aims to characterise *Mansonella spp*. identified during a study assessing the prevalence of lymphatic filariasis in some districts in Ghana.

## Materials and Methods

### Study sites

The study was conducted in 2018 in the East Akyim district (Eastern Region), Adaklu district (Volta Region) and Hohoe district (Volta Region) of Ghana. Communities with M. perstans are shown in Fig 1. In each district participants were recruited from 15 communities based on population proportional to estimated size. These districts experience two raining seasons, with an average rainfall of 513.9 mm to 1099.88 mm. Temperature ranges from 12°C to 32°C. The areas have varying vegetation cover ranging from Forest to savannah grassland. highland areas and in the forest zone which receive the highest rainfall and the dry Sahel-savannah zone in the north(25).

**Fig 1.**
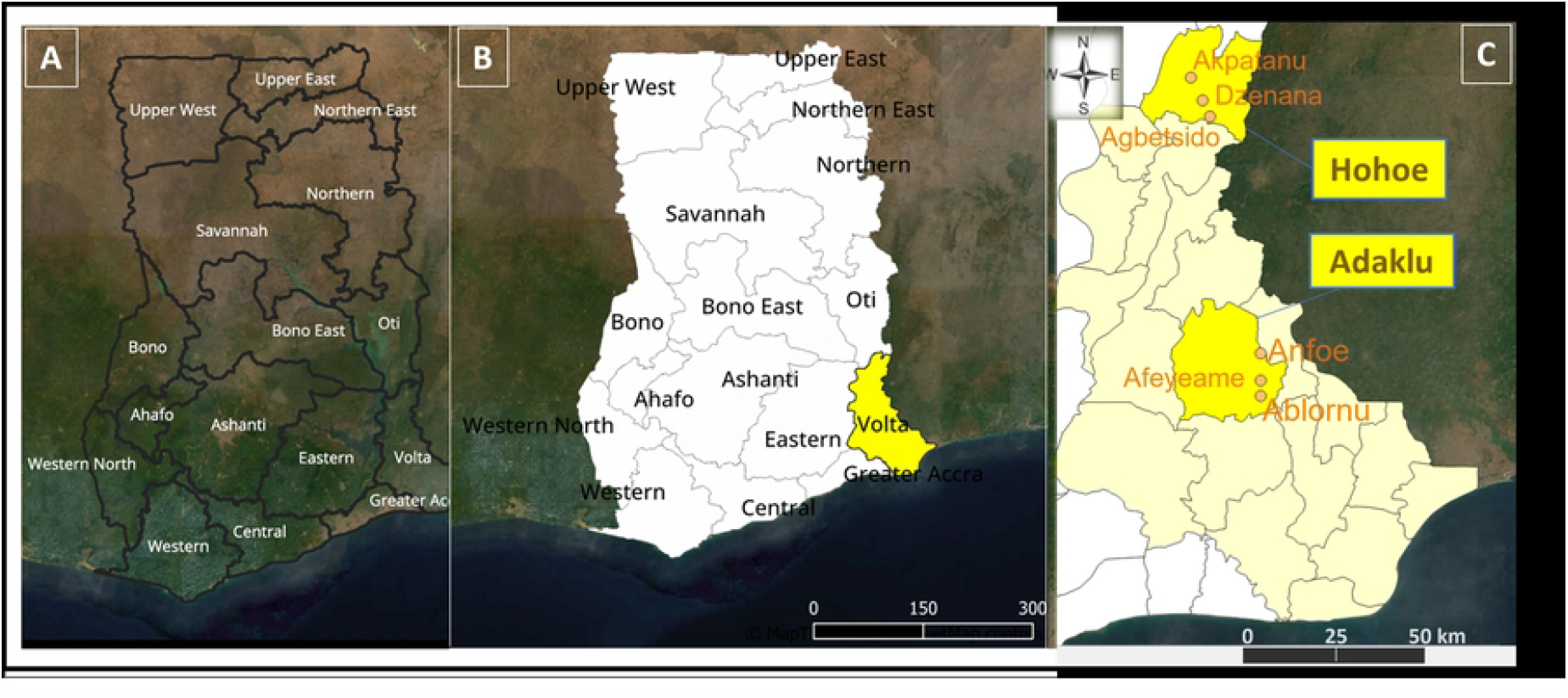
Map of Ghana showing the districts and communities with *M. perstans*. **A**, shows the 16 regions; **B**, shows Volta Region shaded in yellow, and **C**, highlights the two districts, Adaklu and Hohoe with the communities that had study participants infected with *M. perstans*.

### Sample collection

Nighttime blood samples were collected between 21:00 and 2:00 from 50 – 100 randomly selected participants (aged 5 years and above) from each community. 60µl of the blood was used for the preparation of blood films for the identification of *W. bancrofti* microfilariae (mf) and 60 µl to prepare six dry blood spots (DBS), on the six discs of the Tropbio filter paper (Tropbio Pty Ltd, QLD, Australia). The blood films were stained with Giemsa and observed under a compound microscope.

### Sample Processing and Analysis

Following the identification of *M. perstans* in the blood films, the DBS were retrieved and DNA was extracted from 3 discs of the filter paper each per participant. The extractions were done using, the Quick-DNA™ MiniPrep (Zymo Research, USA) according to the manufacturer’s instructions with 75 μL elution volume. PCR was performed to confirm the presence of DNA in the extracted samples, using a human housekeeping gene (CYP11b2). The sample were then screened for Filarial parasites using Nested PCR (26) and CO1 gene (27).

### Detection of Filarial parasites by Nested PCR

Nested PCRs were carried out using four primers described in Tang and colleagues (26). Two primers (FIL-1F; used in the Nest I reaction, and FIL-2R; used in nest II reaction) targeted the 5’-end of the 18S gene. One universal primer (UN1-1R, included in the nest I reaction) targeted 5.8S ribosomal gene. Finally, FIL-2R, a filarial specific reverse primer targeted the ITS-1 region. The nest I reaction, included a FIL-1F (*5’-GTGCTGTAACCATTACCGAAAGG-3’*) and a universal primer, UNI-1R (*5’-CGCAGCTAGCTGCGTTCTTCATCG-3’*)(26) in a total reaction of 15 μL with 300 nM primers, 300 nM dNTPmix, 1X Go-Taq PCR Buffer, 0.033 units of Go-Taq polymerase and 3 μL of the DNA. The PCRs were run at 94°C for 3 min, 35 cycles of 94°C for 30 sec, 60°C for 30 sec and 72°C for 30 sec, and 72°C for 7 min. The nest I was followed with nest II using primers, FIL-2F (5’-GGTGAACCTGCGGAAGGATC-3’), FIL-2R (5’-TGCTTATTAAGTCTACTTAA-3’) in a 15 μL reaction containing, 300 nM primers, 300 nM dNTPmix, 1X Go-Taq PCR Buffer, 0.033 units of Go-Taq polymerase and 2 μL of PCR product from the nest I reaction. The PCRs were run at 94°C for 3 min, 35 cycles of 94°C for 30 sec, 50°C for 30 sec and 72°C for 30 sec, and 72°C for 7 min.

### Gel Electrophoresis and Purification

The Nest II PCR products were run on a 2% agarose gel for 40 min and visualised under blue led light. The bands were excised for further purification using GF-1 Gel DNA recovery kit. The recovered DNA (5μL) were run on a 2% agarose gel to confirm if DNA has been successfully extracted. In total, 30 DNA samples (including a no template control) were sent to Inqaba Biotechnologies, South Africa for Sanger sequencing.

### Sequence Analysis

Sequencing was done in 2021, with two reads obtained per sample. Raw reads were imported into Geneious Prime and poor reads were trimmed with an error probability limit of 0.05 (where more than 5% chance of error per base called are trimmed). The trimmed reads have been deposited in GenBank repository with Accession No. OR488627 to 51. The De Novo Assemble function was then used to generate consensus sequence for each pair rooted with DQ995497 (*Loa loa*). The trimmed consensus sequences were exported to MEGAX(28) to infer ancestry of the sequences. Briefly, the sequences were aligned with MUSCLE and gap open set at -400 and maximum iteration set at 16. Model selection for Maximum Likelihood and the best model estimate of the relation selected was T92+G (based on least Bayesian information criteria (BIC) and corrected Akaike information Criteria (AICc) 1546.049 and 1038.347 respectively (see S4 Table). The rate variation among sites was modelled with a gamma distribution (shape parameter = 0.58). The evolutionary history was inferred by using the Maximum Likelihood method and Tamura 3-parameter model with 500 replicates. A discrete Gamma distribution was used to model evolutionary rate differences among sites (4 categories (+G, parameter = 0.1913)). An analysis of nucleotide variance was generated using popart v1.7 software(29).

### Ethics

Written consent was obtained from all study participants. For children, written parental consent was obtained, and written assent from children 12 to 17 years old. The study was conducted in agreement with the International Council for Harmonization of Technical Requirements for Pharmaceuticals for Human Use (ICH) guidelines on Good Clinical Practice (GCP). Approval for the study was obtained from the Institutional Review Board of the Noguchi Memorial Institute for Medical Research (CPN 061/16-17) and the Ethics Review Committee of the Ghana Health Service (GHS-ERC 06/08/16).

## RESULTS

*M. perstans* was only identified in Hohoe and Adaklu districts (both in the Volta Region). The examination of blood slides revealed *M. perstans* infection in 33/1317 (2.51%) and 22/1355 (1.62%) individuals, respectively in these districts. The mf counts ranged from 16.7 – 12,350.0 mf/ml with a geometric mean of 188.6 mf/ml (S3 Table). None of the people screened in the East Akyim Districts were positive for *Mansonella* parasites.

Thirty-nine (39) DBS from *M. perstans* positive individuals, distributed in 5 communities (Ablornu, Afeyeame, Abledzie, Akpatanu and Dzenana), were retrieved for molecular analysis. Supplementary Fig 1. shows gel electrograms of PCR products with 312 bp (for ITS-1). Overall, 29 samples were sequenced using Sanger sequencing out of which 25 sequences (submitted to GenBank with Accession numbers: OR488627 - OR488651) were obtained for further analysis.

### Phylogenetics

An alignment using MUSCLE in MEGAX was performed with 14 reference sequences (LT623911.1, MZ285895, KR080185.1, EU272180 (*M. ozzardi*), DQ995497 (*Loa loa*), EU272181.1, EU272177.1, KR080189.1, KR080188.1, DQ995498.1, KJ631373.1, KR080187, EU272183.1 EU272182.1) as shown in Fig 2. Following the alignment, the ends were trimmed, and 194 nucleotide sites kept for further analysis. The alignment file can be accessed via: https://github.com/Millicen/Mansonella-perstans/blob/main/Alignment_Fig2.mas. Phylogenetic analysis of the sequences revealed similarity between the Ghana samples and *M. perstans* reference sequences as shown on the tree with 84% and 90% support for the branch lengths (Fig 3). KR080185.1 and MZ285895 which are reference sequences from Gabon and Senegal respectively show differences at some nucleotide positions. We did not see any population structuring in the samples obtained from the different locations in Ghana. Muscle Sequence alignment in MEGA X. Generated with 24 Ghana samples and 14 reference sequences from Africa and Brazil.

**Fig 2.**
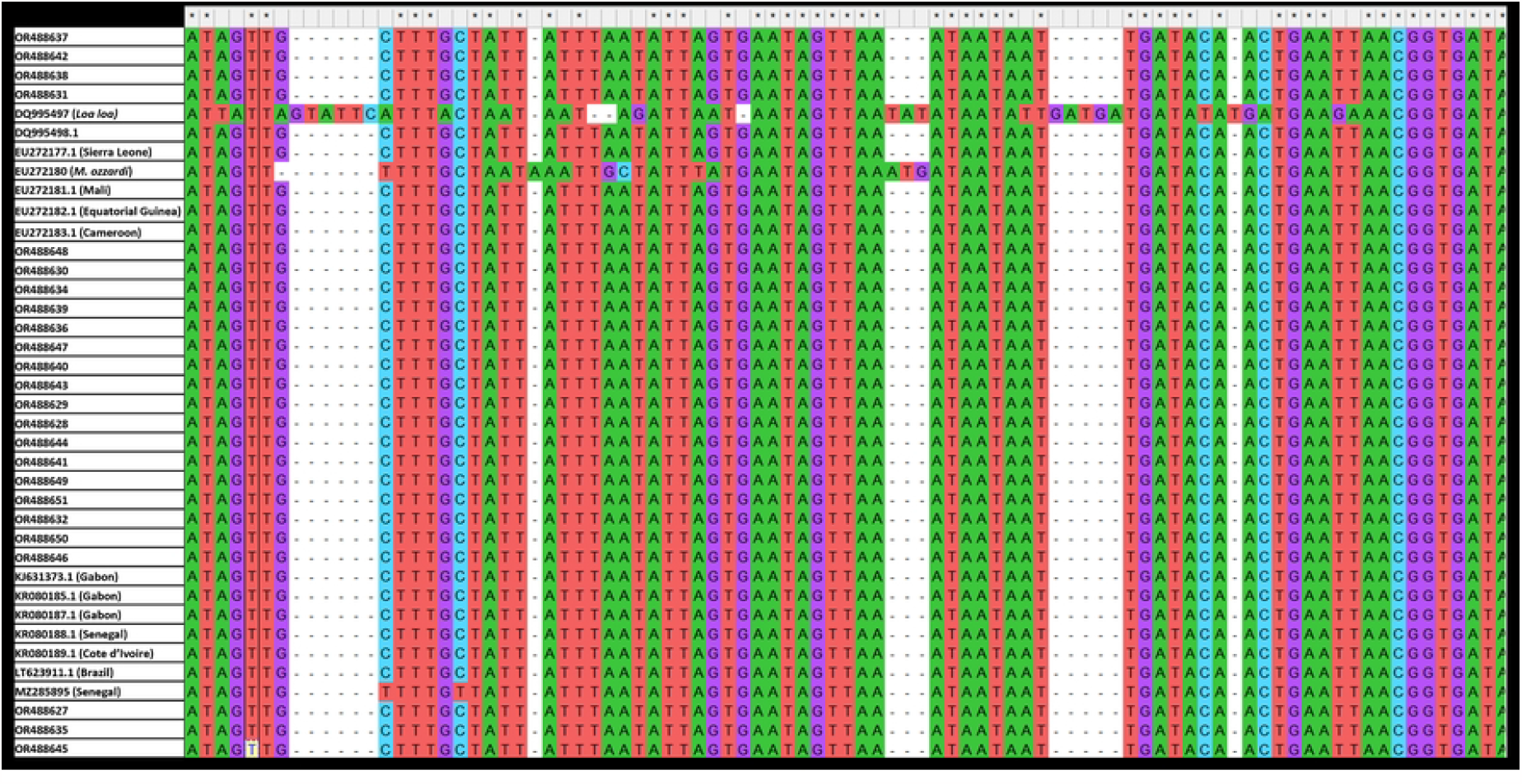
Muscle Sequence alignment in MEGA X. Generated with 24 of the newly sequenced samples and 14 reference sequences from Africa and Brazil. A total of 194 nucleotide sites were retained in the alignment.

**Fig 3.**
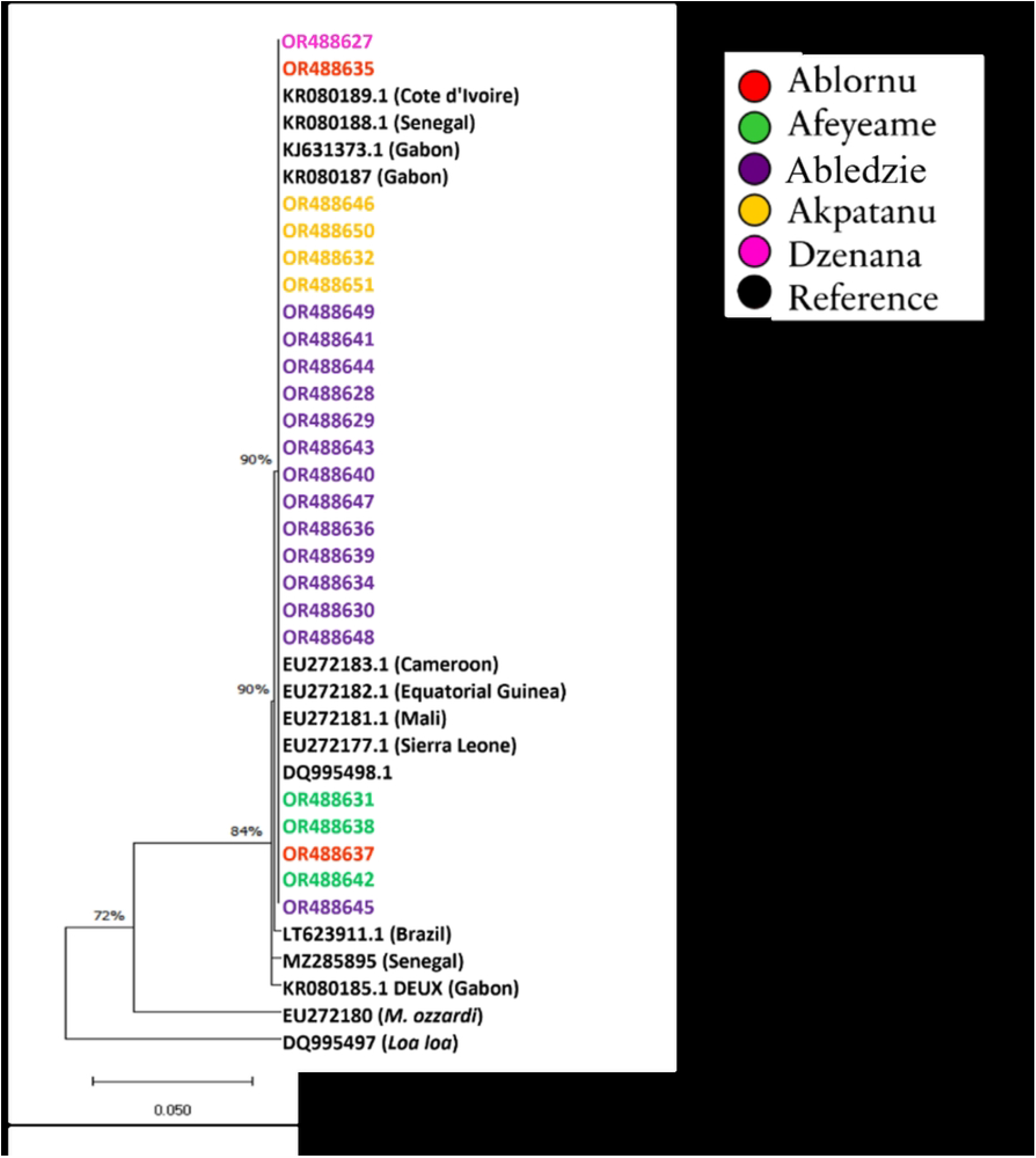
A maximum likelihood tree generated in MEGAX. Varius reference *M. perstans* sequences from Africa (including, Gabon, Senegal, Mali, Sierra Leone, Cote d’Ivoire, Equatorial Guinea and Cameroon) and Brazil, *M. ozzardi* were included with DQ995497 (*Loa loa*) as outgroup. The Ghana samples (OR488627 to 51) are coloured by location. The tree excluded OR488633 due to it short length.

### Haplotype network

An analysis of molecular variance was computed using 24 of the sequenced samples (OR488633 was excluded due to the short length) references, an outgroup (DQ995497) and five *M. perstans* of which two were from Gabon (KR080185.1 and KJ631373.1), one from the Amazona State in Brazil (LT623911.1), one from Senegal (MZ285895) and one from Sierra Leone (EU272177.1) see Fig 4. The nucleotide diversity index was π=0.00898. A significantly high variation (98.24%) among populations, but low variation (1.76%) within populations, were observed with a total variance of 11.89% (fixation index (Phi_ST) = 0.982, p = 0.051). The observed variation was based on 22 bases where, one of the references from Gabon (KJ631373.1) and that of Senegal differed at one and two nucleotide bases respectively, whereas the outgroup had 19 base differences in respect to the Ghana samples. Also, little to no variations were observed between the KJ631373.1 (Gabon), LT623911.1(Brazil) and EU272177.1 (Sierra Leone) and the Ghana samples. No variations were seen in the Ghana samples (p < 0.001).

**Fig 4.**
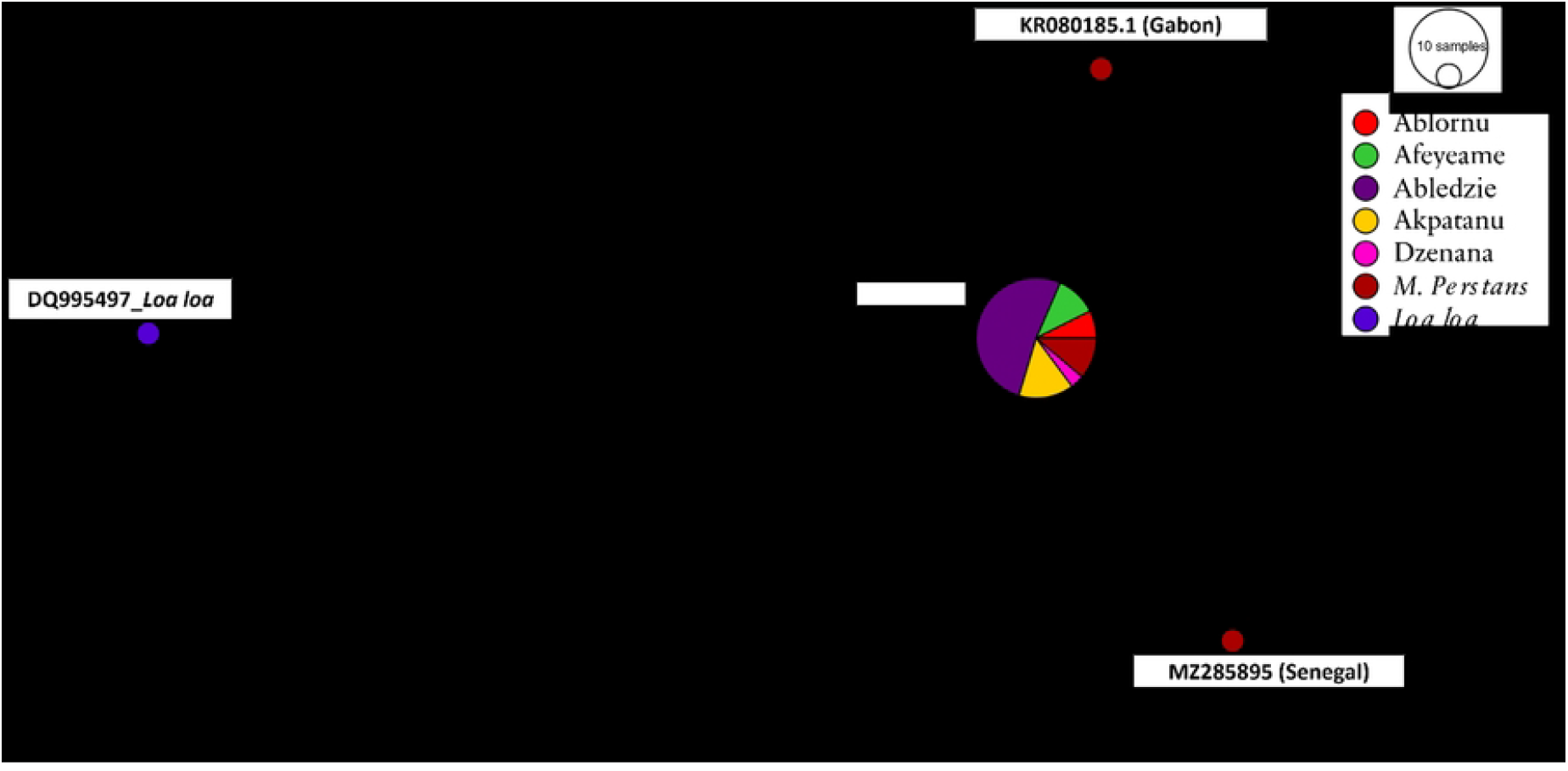
Haplotype network showing nucleotide diversity. Samples are coloured by location. Red and blue nodes indicate reference sequences of *M. perstans* and *Loa loa* respectively and were obtained from the NCBI.

## Discussion

Mansonellosis is rarely symptomatic. However, it may present with itching, joint pains, enlarged lymph glands, and abdominal manifestations. The vague presentations of disease could potentially impact disease outcomes for malaria, tuberculosis, and HIV. Of the three species, *M. perstans* is prevalent in mostly Central and West Africa, and in Central and South America. Although prevalent and co-endemic with other filarial parasites, it is scarcely studied and often picked up by chance. Similarly in this study *M. perstans* were found in blood smears of individuals during lymphatic filariasis studies. Despite the growing evidence of *M. perstans* presence and possible contribution to filarial morbidities, it still remains understudied (17).

In this study we present the first characterisation of *M. perstans* internal transcribed spacer 1 (ITS-1) gene in Ghana. The Ghana samples showed no variations per the phylogenetic analysis suggesting no geographical isolation in the population of *M. perstans* and hence maybe considered as one transmission zone. However, given the small sample sizes with low numbers of representation per site, we are unable to draw firm conclusions on the populations. Many sources of *M. perstans* infection have been linked to Africa. In Italy and Spain, *M. perstans* were identified in immigrants from sub-Saharan Africa (30). In the Caribbean and the Americas, infections have been linked to the trans-Atlantic slave trade (31). The clustering of the Ghana samples and other sequences from Gabon and Sierra Leone with the *M. perstans* ITS-1 (LT623911.1) from Brazil Amazona State and other countries in Africa supports work by Tavares da Silva and colleagues (31) that showed grouping of sequences from Cameroon, Côte d’Ivoire, Equatorial Guinea, Gabon, Mali, and Sierra Leone to LT623911 (from Brazil). Their work also supported the origin of *M. perstans* from Africa into Latin America following divergence from *M. perstans* “deux” clade from Gabon (31).

While the identification of *M. perstans* in this study was conducted based on the ITS-1, a newer method based on the *M. perstans* Repeat 1 (*Mp*R1) has been developed (8). The recent publication of the *M. perstans* genome (32) will also help further studies and understanding of this very neglected parasite.

This report of *M. perstans* in the Hohoe and Adaklu districts, in the East of Ghana, effectively indicates the presence of the infection in the lower half of the country. The distribution of the infection within these districts and coexistence with other filarial infections presents opportunities for the integration of Mansonellosis surveillance with existing LF and onchocerciasis programs.

## Conclusion

Despite the potential impact of *M. perstans* on diseases of public health importance, it receives little attention. Consequently, there is no standard treatment and surveillance to track the extent of endemicity. Although symptoms are not clearly defined it is believed to impact host immune reactions and may play a crucial role in disease pathogenesis especially where it is co-endemic with other filarial parasites. In this study, we present the first characterisation of *M. perstans* in Ghana. Furthermore, the phylogenetics analysis reveals clustering of isolates from Ghana with that of Brazil. We did not observe any differences in the sequences obtained from the study sites which could probably be due to small sample size. We therefore advocate for population-based studies with a bigger target (or whole genome sequence data) that will include more samples from different locations across different geographical settings. Furthermore, the coexistence of *M. perstans* with other blood filarial worms presents a unique opportunity for screening one sample from a single individual allowing for parallel monitoring as well as assessing the public health impact of mansonellosis in populations.

## Acknowledgement

Dr. Jewelna Akorli provided access to Geneious Software for data analysis. Dr. Shannon Hedtke provided guidance on data analysis.

## Supporting information

**S1 Fig 1. Gel electrogram showing PCR products of samples after Nested reaction**. Where Lanes 1 to 9 = samples; +ve = positive control; - = Negative control

**S2 Table. GPS coordinates of the communities from Adaklu and Hohoe districts**.

**S3 Table. Number of samples that were positive for *M. perstans* following microscopy**. Those that were submitted for Sanger sequencing are indicated with *. The highest mf load (12350.0 mf/mL) was recorded in Abledzie.

**S4 Table. Models estimation and selection inferring relationships** T92+G model was selected to build the tree based on low BIC and AICc values.

